# Sleep loss disrupts hippocampal memory consolidation via an acetylcholine- and somatostatin interneuron-mediated inhibitory gate

**DOI:** 10.1101/2020.08.02.233080

**Authors:** James Delorme, Femke Roig Kuhn, Lijing Wang, Varna Kodoth, Jingqun Ma, Sha Jiang, Sara J. Aton

## Abstract

Sleep loss profoundly disrupts consolidation of hippocampus-dependent memory. To better characterize effects of learning and sleep loss on the hippocampal circuit, we quantified activity-dependent phosphorylation of ribosomal subunit S6 (pS6) across the dorsal hippocampus of mice. We find that pS6 in enhanced in the dentate gyrus (DG) following single-trial contextual fear conditioning (CFC), but is reduced throughout the hippocampus after brief sleep deprivation (SD) – a manipulation which disrupts contextual fear memory (CFM) consolidation. To characterize cell populations with activity affected by SD, we used translating ribosome affinity purification (TRAP)-seq to identify cell type-specific transcripts on pS6 ribosomes after SD vs. sleep. Cell type-specific enrichment analysis (CSEA) of these transcripts revealed that hippocampal somatostatin-expressing (Sst+) interneurons, and cholinergic and orexinergic inputs to hippocampus, are selectively activated after SD. We used TRAP targeted to hippocampal Sst+ interneurons to identify cellular mechanisms mediating SD-driven Sst+ interneuron activation. We next used pharmacogenetics to mimic the effects of SD, selectively activating hippocampal Sst+ interneurons while mice slept in the hours following CFC. We find that activation of Sst+ interneurons is sufficient to disrupt CFM consolidation, by gating activity in surrounding pyramidal neurons. Pharmacogenetic inhibition of cholinergic input to hippocampus from the medial septum (MS) promoted CFM consolidation and disinhibited neurons in the DG, increasing pS6 expression. This suggests that state-dependent gating of DG activity is mediated by cholinergic input during SD. Together these data provide evidence for an inhibitory gate on hippocampal information processing, which is activated by sleep loss.

## Introduction

Hippocampal plasticity and memory storage are gated by vigilance states. In both human subjects and animal models, sleep loss disrupts consolidation of multiple types of hippocampus-dependent memories (Abel et al., 2013; Havekes and Abel, 2017). This effect has been extensively studied in mice, where as little as a few hours of experimental sleep deprivation (SD) can disrupt hippocampally-mediated consolidation of object-place memory (Havekes et al., 2016; Prince et al., 2014; Vecsey et al., 2009) and contextual fear memory (CFM) (Graves et al., 2003; Ognjanovski et al., 2018). Recent work has characterized biochemical pathways involved in memory consolidation which are disrupted in the hippocampus by SD (Aton et al., 2009b; Havekes et al., 2016; Tudor et al., 2016; Vecsey et al., 2009). However, much less is known about how SD affects hippocampal microcircuit function.

SD disrupts patterns of hippocampal network activity which are associated with memory consolidation. For example, SD interferes with network activity changes induced in hippocampal area CA1 by prior learning (CFC), which predict successful CFM consolidation (Ognjanovski et al., 2018; Ognjanovski et al., 2014). The reason for this SD-mediated disruption is unknown. Recently, activity-dependent regulation of protein translation machinery within dorsal hippocampus was found to be essential for sleep-dependent memory consolidation (Tudor et al., 2016). SD interferes with biochemical pathways which drive increased protein synthesis following learning (Tudor et al., 2016; Vecsey et al., 2012). This suggest a link between state-dependent changes in network activity and biosynthetic events necessary for appropriate memory consolidation.

To better characterize the link between neuronal activity and protein synthesis in the hippocampus during CFM consolidation, we characterized effects of learning and subsequent sleep or SD on activity-dependent phosphorylation of ribosomal subunit S6 (pS6). We find that CFC increases pS6 phosphorylation at a terminal serine residue (pS6 Ser242-244), and that SD reduces pS6 Ser242-244 throughout the dorsal hippocampus. To identify cell populations differentially expressing pS6 after sleep vs. SD, we used a pSer242-244 as an affinity tag for translating ribosome affinity purification (pS6-TRAP). We then identified modules of cell-type specific transcripts with expression correlated to wake time in sleeping and SD mice, and verified these findings with qPCR. These analyses indicate that SD selectively activates (i.e., leads to increased pS6 expression in) hippocampal Sst+ interneurons, and orexinergic (lateral hypothalamic) and cholinergic (MS) neurons which send input to the hippocampus. We used TRAP in Sst+ interneurons (Sst-TRAP) to verify that activity-dependent transcripts are increased in these neurons with SD. To assess how increased activity in the hippocampus Sst+ interneuron population during SD affects memory consolidation, we used pharmacogenetics to selectively activate these neurons in the hours following CFC. We find the mimicking the effects of SD on Sst+ interneuron activity is sufficient for disruption of CFM consolidation in freely-sleeping mice. Lastly, we tested the hypothesis that state-dependent regulation of the dorsal hippocampal network is mediated by changes in activity of MS cholinergic neurons. We find that pharmacogenetic inhibition of cholinergic input to hippocampus following CFC promotes CFM consolidation, and increases pS6 expression in dorsal hippocampus. Together, these data provide evidence for a state-dependent gate on network activity in the hippocampus, regulated by Sst+ interneurons and MS cholinergic input, which causes SD-induced disruption of memory consolidation.

## Results

### Learning increases and sleep loss decreases phosphorylation of S6 in the hippocampus

Since brief sleep deprivation (SD) of only a few hours is sufficient to disrupt many forms of hippocampus-dependent memory consolidation in mice (Graves et al., 2003; Havekes et al., 2016; Ognjanovski et al., 2018; Prince et al., 2014; Vecsey et al., 2009), we first characterized the effects of 3-h SD on S6 phosphorylation. Following 5 days of habituation to handling, beginning at lights on (ZT0), mice either had continuous SD by gentle handling or were allowed *ad lib* sleep (Sleep) prior to sacrifice at ZT3 (**Figure 1A**). Because S6 is sequentially phosphorylated at five serine residues, we first quantified phosphorylation using an antibody recognizing the initial Ser235-236 phosphorylation sites (pS6 235-236). Consistent with previous reports (Tudor et al., 2016), 3-h SD did not alter either the number of pS6(Ser235-236)+ neurons in the dentate gyrus (DG) or the intensity of pS6 235-236 staining in the pyramidal cell body layers of CA1 or CA3 (**Figure 1C**). We then quantified phosphorylation at the terminal S6 sites (Ser244-247; hereafter referred to simply as pS6). in SD mice, we observed a significant decrease in the number of pS6+ neurons in the dentate gyrus, and reduced intensity of pS6+ staining in pyramidal cell body layers of CA1 and CA3 (**Figure 1B, C**). In contrast, neocortical regions adjacent the dorsal hippocampus (i.e., primary somatosensory cortex) showed increased numbers of pS6+ neurons at both sites after 3-h SD (**Figure S2**).

**Figure 1.**
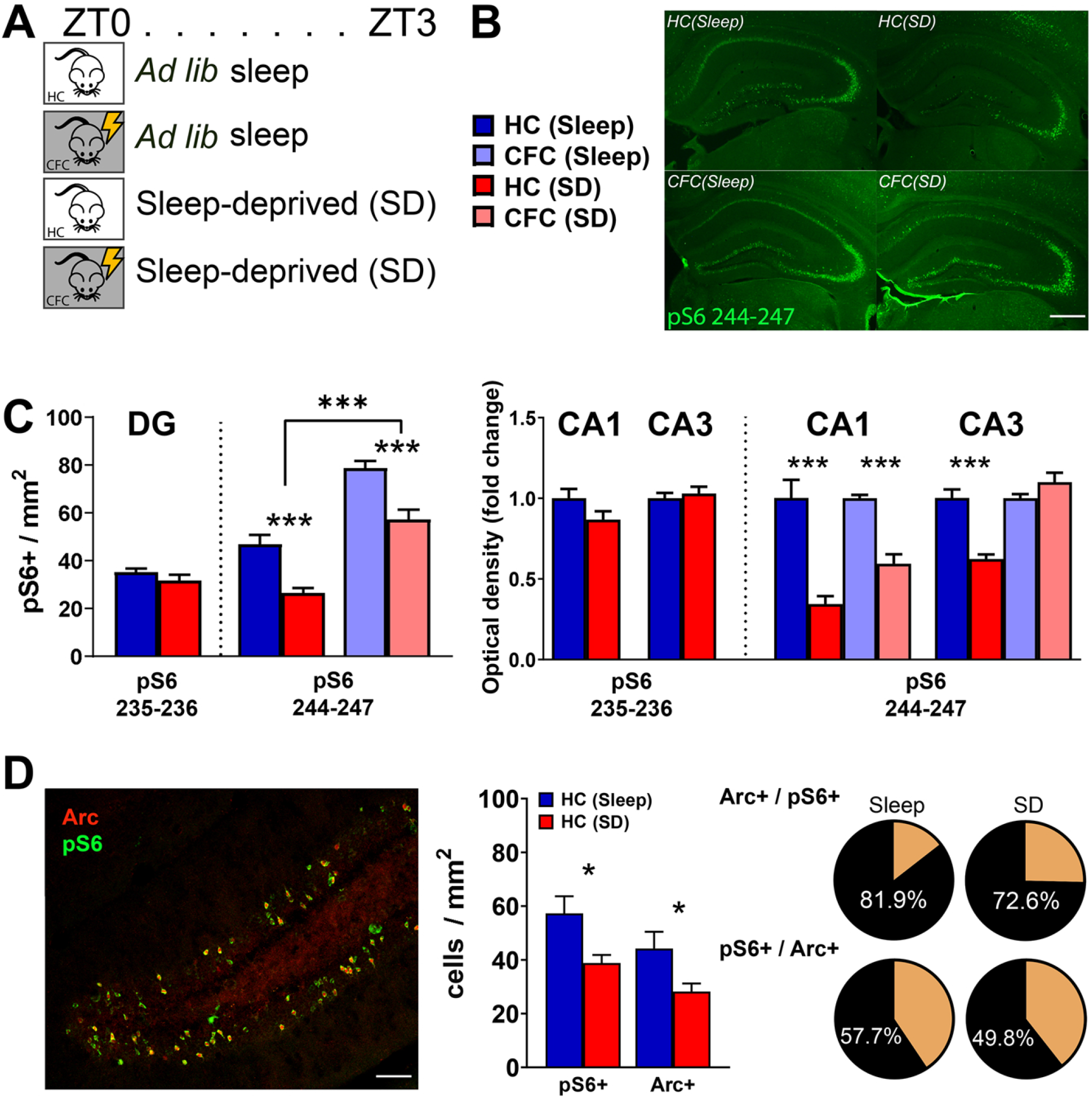
Hippocampal S6 phosphorylation increases after learning and is reduced by sleep deprivation (SD). ***A)*** Experimental paradigm. Mice underwent single-trial contextual fear conditioning (CFC) at ZT0, or were left in their home cage (HC). Over the next 3 h, mice in CFC and HC groups were then permitted *ad lib* sleep (Sleep) or were sleep-deprived (SD) by gentle handling. ***B)*** Fluorescent images of pS6 (S244-247) staining in the dorsal hippocampus of representative mice in the four treatment groups. Scale bar = 500 μm. ***C) Left:*** pS6+ neurons in DG were counted using antibodies detecting S6 phosphorylation at either S235-236 or S244-247 sites. SD selectively reduced S244-247 pS6+ neurons in both HC (*n* = 5/group) and CFC (*n* = 5/group) mice. CFC increased the number of pS6+ neurons (two-way ANOVA: main effect of CFC, *F* = 87.09, *p* < 0.001; main effect of SD, *F* = 38.94, *p* < 0.001; CFC × SD interaction, *N.S.*). ***Right:*** pS6 expression in pyramidal cell layers CA1/CA3 was quantified as background subtracted optical density. SD values were calculated as the fold change relative to the Sleep condition HC + Sleep or CFC + Sleep, respectively. pS6 (S244-247) OD was reduced in both CA1 (*p* < 0.001, Student’s t-test) and CA3 (*p* < 0.001) after SD in HC mice. After CFC, SD reduced pS6 expression in CA1 neurons (*p* < 0.001) following SD. ***D) Left:*** Representative image of pS6 and Arc colocalization in DG. Scale bar = 500 μm. ***Middle:*** Quantification of pS6 and Arc expression in HC + Sleep and HC + SD mice (*n* = 5/group). 3-h SD reduced both Arc+ (*p* < 0.05, Student’s t-test) and pS6+ (*p* < 0.05) neurons in DG. ***Right:*** ~77% of Arc+ DG neurons expressed pS6+; ~54% of pS6+ neurons expressed Arc.

We verified that S6 phosphorylation is neuronal activity-driven, consistent with previous reports (Pirbhoy et al., 2016), by quantifying co-expression of the activity-regulated protein Arc in pS6+ neurons. Consistent with our previous findings (Delorme et al., 2019), 3-h SD reduced numbers of both Arc+ and pS6+ neurons in the DG. Arc and pS6 were co-localized to a similar extent in DG of both Sleep and SD mice (**Figure 1D**), with 77 ± 3.2% of Arc+ DG neurons also being pS6+, and 54 ± 2.8% of pS6+ neurons also being Arc+ (mean ± SEM from *n* = 10 mice).

We next tested whether hippocampal S6 phosphorylation was affected by learning a hippocampus-dependent memory task. Mice underwent single-trial contextual fear conditioning (CFC; in which exploration of a novel chamber is paired with a foot shock) or, for comparison, were left in their home cage (HC) at lights on. After this, both CFC and HC mice were either allowed *ad lib* sleep, or had SD by gentle handling in their home cage. In freely-sleeping mice, CFC increased the number of pS6+ DG neurons at both 30 min and 3 h post-CFC, relative to HC controls (two-way ANOVA: main effect of time, *F* = 50.63, *p* < 0.001; main effect of learning, *F* = 33.59, *p* < 0.001; time × learning interaction, *F* = 2.22, *p* = 0.16) (**Figure S1**). In contrast, CFC did not alter pS6 expression CA1 or CA3 of freely-sleeping mice, relative to HC controls. Consistent with greater pS6 expression in the hippocampus after periods rich in sleep, pS6+ neurons increased between the two timepoints (ZT0 vs. ZT3) in both HC + Sleep and CFC + Sleep mice (**Figure S1**). 3-h SD disrupted pS6 expression in the hippocampus following CFC, with fewer pS6+ neurons in the DG and reduced pS6+ expression in CA1 (**Figure 1C**). Taken together, these data suggest that learning increases and SD reduces S6 phosphorylation in the hippocampus.

### Identification of hippocampal cell types with altered S6 phosphorylation during SD

We next used an unbiased RNA-seq approach to identify cells in which pS6 expression differs between Sleep and SD. Using pS6 as an affinity tag to isolate ribosomes and associated transcripts in active cells, we performed pS6 translating ribosome affinity purification (pS6-TRAP) (Knight et al., 2012). Hippocampi were collected from CFC and HC mice after 3 h *ad lib* sleep or SD. Ribosome-associated transcripts were then isolated by pS6-TRAP for RNA-seq. To identify clusters of co-regulated transcripts in our RNA-seq data (such as might be expected for genetically-defined cell types), we used weighted gene correlation network analysis (WGCNA) (Langfelder and Horvath, 2008) on transcripts with a variance greater than 0.03 (*n* = 1662 transcripts). WCGNA yielded 10 clusters (modules) of highly correlated transcripts in our data, and a separate (Gray) cluster representing unassigned (uncorrelated) transcripts (**Figure 2A, Supplemental Table 1**). To determine which modules’ expression varied as a function of Sleep vs. SD, we correlated the level of expression of module eigengenes with the percent of time mice spent awake over the 3 h prior to sacrifice (Sleep = 25.6 ± 2.2%, SD = 100 ± 0.0% [mean ± SEM]). Results from the analysis revealed two significantly correlated eigengene clusters (Brown, Magenta) whose expression negatively correlated with sleep time (**Figure 2A**). Since these represented sub-clusters of the same module, we combined them for further analysis (Brown/Magenta cluster).

**Figure 2.**
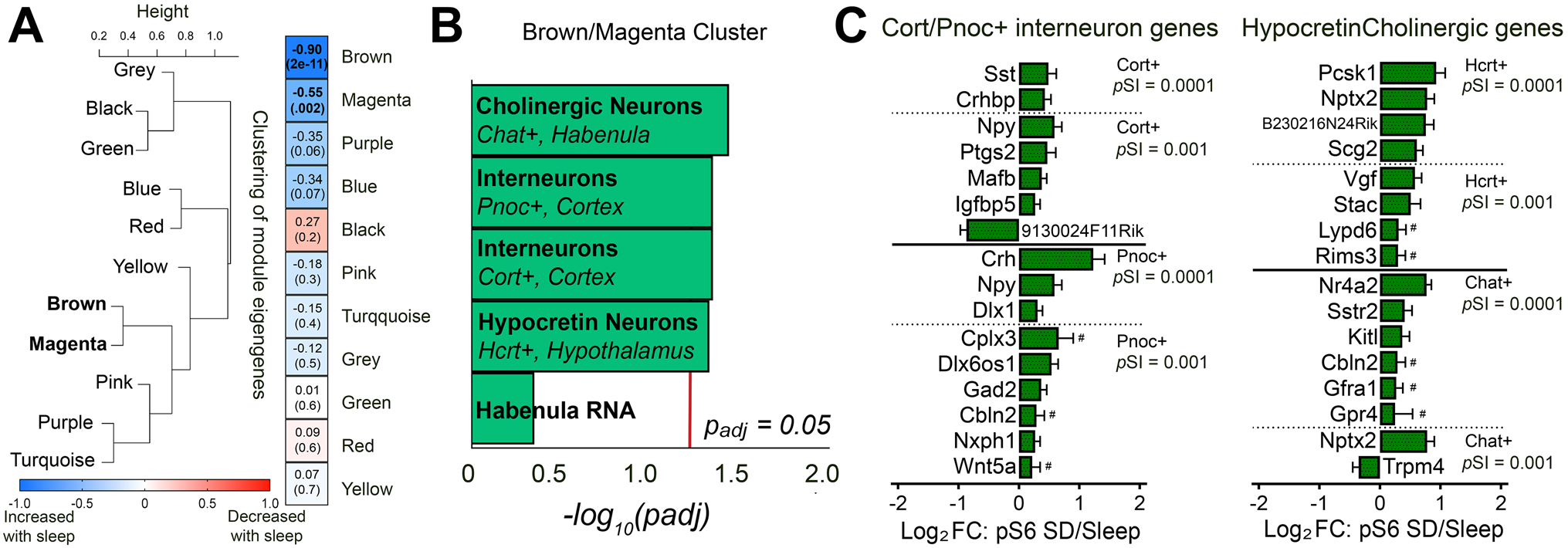
Phosphorylated ribosome capture following SD selectively enriches transcripts specific to GABAergic, cholinergic, and orexinergic neurons. ***A) Left:*** Weighted gene co-expression analysis (WGCNA) identified modules of similarly correlated pS6 transcripts. Each module is identified with a color name, Grey represents transcripts not assigned to a co-expression module. ***Right:*** Correlation between eigengene expression in each module and total sleep time prior to sacrifice (*R*- and *p*-values for Pearson correlation in parentheses for each module). ***B)*** Transcripts in the Brown/Magenta module with expression correlated to sleep time were used for cell type-specific expression analysis (CSEA). The Brown/Magenta transcripts showed significant overlap with mRNAs enriched most selectively (*p*SI < 0.0001) in cholinergic (Epi.ChAT, *p*_adj_ = 0.036), orexinergic (Hyp.Hcrt, *p*_adj_ = 0.047), and GABAergic (Ctx.Pnoc, *p*_adj_ = 0.045; Ctx.Cort, *p*_adj_ = 0.045) neuron populations. ***C)*** Deseq2 Log_2_FC (SD/Sleep) values from Brown/Magenta transcripts identified by CSEA. All genes are statistically significant (*p*_adj_ < 0.1) unless otherwise indicated (# indicates *p*_adj_ > 0.1).

Since our data suggested that the population of pS6+ neurons in the hippocampus may differ in freely-sleeping and SD mice (**Figure 1**), we used cell type-specific expression analysis (CSEA) (Xu et al., 2014) to quantify cell type-specifying transcripts represented in the Brown/Magenta cluster, which were significantly affected by SD. CSEA was used to generate a *p*_adj_ value for overlap between transcripts in Brown/Magenta cluster and known cell type-specific enriched transcripts of a particular specificity index *p* value (*p*SI) (based on a multiple comparisons-corrected Fisher’s exact test). Using the most stringent CSEA (*p*SI < 0.0001), we identified Brown/Magenta cluster transcripts as mRNAs expressed most selectively in cholinergic (Chat+) neurons (*p*_adj_ = 0.036), orexinergic (Hcrt+) neurons (*p*_adj_ = 0.047), and Pnoc+ and Cort+ interneurons (*p*_adj_ = 0.045) (**Figure 2B-C, Supplemental Table 1**). This suggests that after SD, pS6 is associated with more transcripts from hippocampal neurons similar to these neuron types, despite the fact that overall pS6 expression is reduced after SD. The former likely reflects transcripts present in orexinergic inputs to the hippocampus from lateral hypothalamus and cholinergic input from the medial septum, respectively - both of which are more active during active wake vs. sleep (Kiyashchenko et al., 2002; Teles-Grilo Ruivo et al., 2017). With respect to the latter finding, overlap between the Brown/Magenta cluster and transcripts expressed selectively in Cort+ and Pnoc+ interneurons (Doyle et al., 2008; Taniguchi et al., 2011) included transcripts encoding interneuron-specific transcription factors (*Dlx1*) and secreted neuropeptides somatostatin (*Sst*), neuropeptide Y (*Npy*), and corticotrophin-releasing hormone (*Crh*).

### Hippocampal somatostatin-expressing (Sst+) interneurons and cholinergic inputs show increased activity during brief SD

Sst and Npy neuropeptides are co-expressed in dendritic-targeting interneurons in DG, CA3, and CA1, and play a role in gating neighboring neuronal activity (Kosaka et al., 1998; Pelkey et al., 2017; Stefanelli et al., 2016). To confirm enrichment of *Sst, Npy,* and other CSEA-identified transcripts in the pS6+ cell population after SD, we carried out a second experiment in which CFC and HC mice either were allowed 5 h of *ad lib* sleep or underwent 5-h SD. pS6-TRAP was followed by quantitative PCR (qPCR) to measure cell type-specific transcripts from the hippocampus. We found that independent of prior training (CFC or HC), SD caused similar enrichment for transcripts present in GABAergic neurons in pS6-TRAP. While *Gad67* and *Pvalb* transcripts were only moderately increased following 5-h SD, *Sst* and *Npy* showed large increases (**Figure 3A**). SD also increased *Cht* expression in both CFC and HC mice. These data support our unbiased CSEA-based finding of increased abundance of Sst-expressing (Sst+) interneuron and cholinergic neuron markers in the SD pS6+ population.

**Figure 3.**
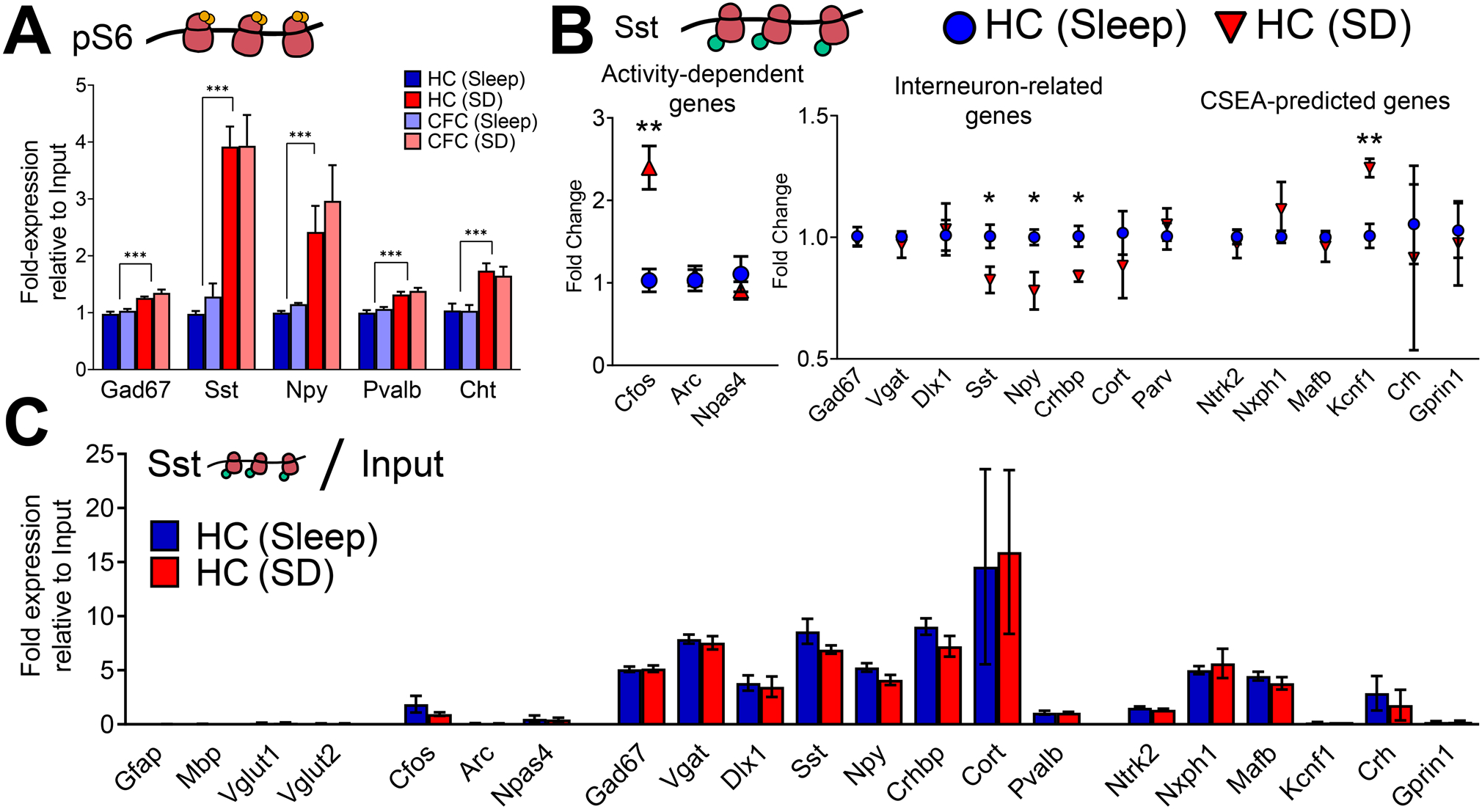
SD increases activity in Sst+ interneurons. ***A)*** qPCR data for pS6-associated transcripts from CFC (*n* = 6/group) or HC (*n* = 5/group) mice with 5 h subsequent *ad lib* sleep or SD (two-way ANOVA: main effect of SD, *p* < 0.001; main effect of CFC, *N.S.*; CFC × SD interaction, *N.S.*). ***B)*** Expression of cell type-specific markers in mRNA from Sst-TRAP vs. Input. SST de-enriched transcripts expressed in glial cells, and preferentially enriched transcripts expressed in Sst+ (GABAergic) interneurons. These enrichment values did not differ between HC+ Sleep (*n* = 5) and HC + SD (*n* = 4) mice. ***C)*** Changes in expression of activity-regulated, interneuron-specific, and CSEA-predicted transcripts associated with Sst+ interneuron ribosomes following 3-h of *ad lib* sleep or SD. Sleep vs. SD, ***, **, and * indicate *p* < 0.001, *p* < 0.01, and *p* < 0.05, respectively, Student’s t-test.

Because these data suggest that despite reduced total activation of hippocampal neurons during SD, Sst+ interneurons in the hippocampus are more activated, we next quantified expression of activity markers in Sst+ interneurons directly. We used TRAP to isolate mRNAs associated with translating ribosomes in this cell population using *SST-IRES-CRE* transgenic mice expressing hemagglutinin (HA)-tagged Rpl-22 (RiboTag) in a Cre-dependent manner (Sanz et al., 2019). Ribosome-associated transcripts from the hippocampus of these mice, isolated following a 3-h period of sleep or SD, were quantified with qPCR. We first verified enrichment of cell type-specific (i.e., Sst+ interneuron-specific) mRNAs, by comparing transcript levels from TRAP vs. Input (whole hippocampus) mRNA. Sst-TRAP significantly de-enriched glial (G*fap, Mbp*) and excitatory neuron (*Vlugt1*, *Vglut2*) selective transcripts, and significantly enriched for Sst+ interneuron-expressed transcripts *Gad1, Vgat, Sst, Npy, and Crhbp* (**Figure 3B**). We then tested whether 3-h SD increased the expression of activity-regulated transcripts in Sst+ interneurons, and found that *Cfos* (but not *Npas4* or *Arc*), was significantly elevated at Sst+ interneurons’ ribosomes after SD (**Figure 3C**). We also tested whether SD-driven increases in *Sst*, *Npy*, and *Crhbp* in pS6-TRAP were due to increased expression levels within Sst+ interneurons. Using qPCR for these neuropeptide transcripts in mRNA isolated using Sst-TRAP pulldown, we found that following 3-h SD, *Sst*, *Npy*, and *Crhbp* transcripts were all *less abundant*, rather than more abundant (**Figure 3C**). The same change was not present in Input (whole hippocampus) mRNA, where expression of *Sst, Npy,* and *Crhbp* mRNAs were all unchanged by SD (**Figure S3**). We also used qPCR to quantify expression of other cell type-specific transcripts identified in pS6-TRAP by WCGNA/CSEA. Of the transcripts tested, we found that SD increased expression of *Kcnf1*, encoding the voltage-gated potassium channel subunit Kv2.1 in Sst+ interneurons (**Figure 3C**). Critically, greater expression of *Kcnf1* is correlated with reduced action potential threshold and increased neuronal firing rate (Bomkamp et al., 2019). Taken together, these data support the conclusion that SD-induced increases in *Sst*, *Npy*, and *Crhbp* transcripts in pS6-TRAP reflect increases in the activity, and thus pS6 expression, within Sst+ interneurons.

To further validate increases in Sst+ interneuron activity after SD, we examined SD-driven changes in pS6 expression in Sst+ and parvalbumin-expressing (Pvalb+) interneurons in the dorsal hippocampus using immunohistochemistry (**Figure 4A**). As observed previously (**Figure 1**), 3-h SD reduced the total number pS6+ neurons in the DG (**Figure 4B**). However, at the same time, 3-h SD increased pS6 expression among Sst+ interneurons in the DG, and showed a strong trend for increased expression in CA3 Sst+ interneurons (**Figure 4C**). Overall numbers of Sst+ interneurons were similar between Sleep and SD mice, and consistent with qPCR results from Sst-TRAP, the intensity of Sst staining among Sst+ interneurons was decreased after SD (**Figure S4**).

**Figure 4.**
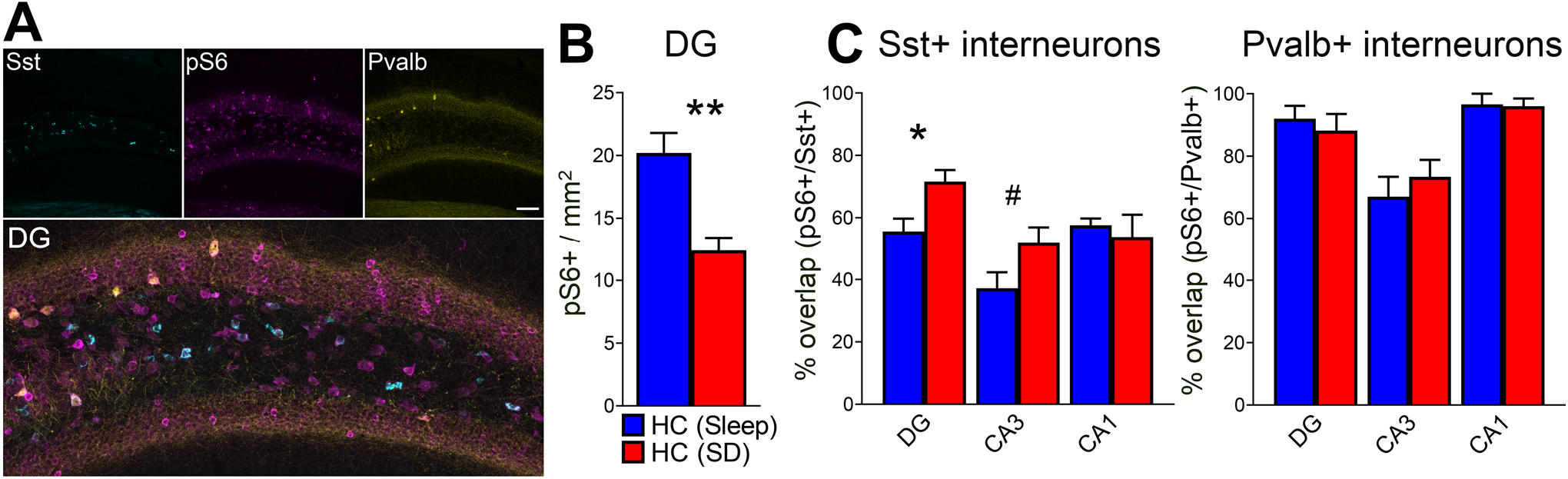
DG Sst+ interneurons show increased pS6 expression following SD. ***A)*** Representative images showing expression of Sst, Pvalb, and pS6 in DG (scale bar = 100 μm). ***B)*** DG pS6+ neurons decreased following 3-h SD (*p* < 0.01, Student’s t-test, *n* = 5 mice/group) ***C)*** pS6 colocalization in Ssy+ and Pvalb+ DG interneurons was compared for HC mice after 3-h SD or *ad lib* sleep. SD elevated pS6 expression in Sst+ interneurons in the DG hilus (*p* < 0.05, Student’s t-test) and trended in CA3 (*p* < 0.1). pS6 expression in Pvalb+ interneurons was unaffected by SD. Sleep vs SD, ***, **, *, and # indicated *p* < 0.001, *p* < 0.01, *p* < 0.05, and p < 0.1 respectively, Student’s t-test.

### Mimicking SD-driven increases in Sst+ interneuron activity in the hippocampus disrupts sleep-dependent memory consolidation

Because the SD-associated increase in Sst+ interneuron activity has the potential to profoundly suppress surrounding hippocampal network activity (Raza et al., 2017; Stefanelli et al., 2016), we next tested how this process affects sleep-dependent memory consolidation. To test this, we transduced the dorsal hippocampus of *SST-IRES-CRE* mice with an AAV vector to express either the activating DREADD hM3Dq-mCherry or mCherry alone in a Cre-dependent manner (**Figure 5A**). To confirm effects of pharmacogenetic manipulation on Sst+ interneuron and surrounding DG granule cells’ neuronal activity, hM3Dq- and mCherry-expressing (*n* = 5 and 4, respectively) mice were injected with clozapine-N-oxide (CNO; 3 mg/kg) at lights on and allowed 3 h *ad lib* sleep in their home cage prior to sacrifice. hM3Dq-mCherry-expressing DG neurons showed a significantly higher level of cFos expression at this time point (hM3Dq: 68.0 ± 15.9% vs. mCherry: 2.0 ± 1.4%; *p* < 0.01 Student’s t-test) **(Figure 5A, Figure 5B)**. To assess effects of CNO on Sst+ interneuron-mediated inhibition in the surrounding DG, we quantified cFos expression in non-transduced neurons in the DG granule cell layer. hM3Dq expression significantly reduced numbers of cFos+ neurons in the surrounding DG relative to mCherry-expressing control mice (*p* < 0.05, Student’s t-test) **(Figure 5C)**. To test how activation of Sst+ interneurons affects memory consolidation, hM3Dq-mCherry- and mCherry-expressing mice underwent single-trial CFC at lights on, after which they were immediately injected with CNO, and returned to their home cage for *ad lib* sleep. 24 h later, at lights on, mice were returned to the CFC context to assess CFM. Mice expressing hM3Dq showed significant decreases in context-specific freezing compared to mCherry-expressing control mice (*p* < 0.001, Student’s t-test). (**Figure 5D**). Together, these data show that sleep-dependent consolidation of CFM can be disrupted via activation of Sst+ interneurons, which provide strong inhibition to the surrounding hippocampal network.

**Figure 5.**
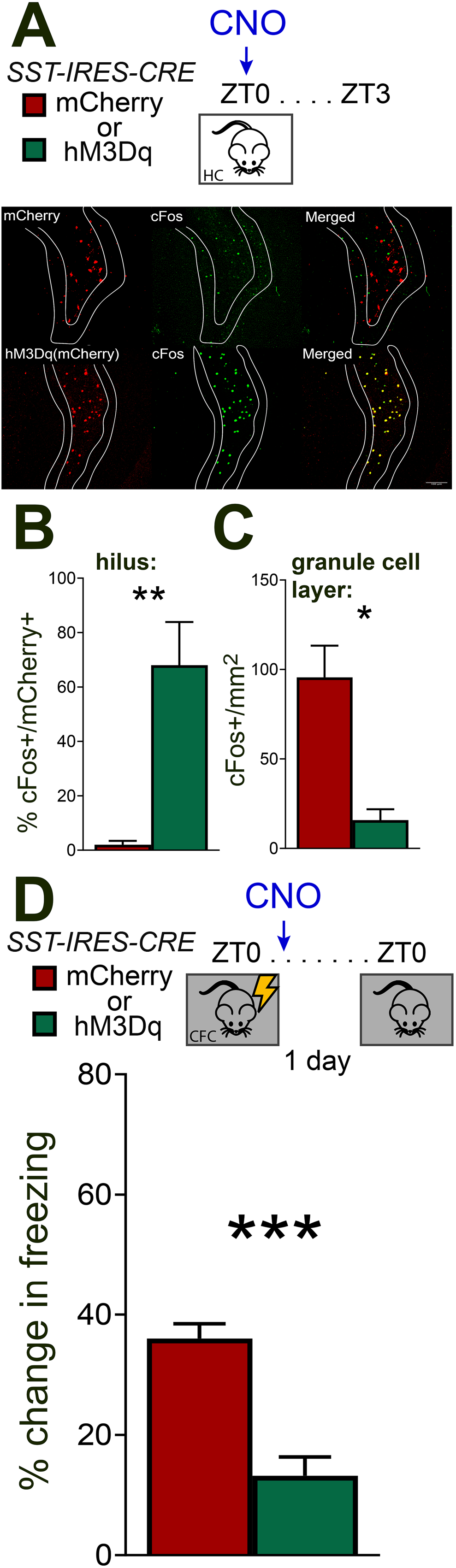
Mimicking SD effects on activity in Sst+ interneurons impairs sleep-dependent memory consolidation. ***A) Top:*** Experimental design: *SST-IRES-CRE* mice expressing either mCherry (*n* = 4) or hM3Dq-mCherry (*n* = 5) in dorsal hippocampus were injected with CNO (3 mg/kg, i.p.) at lights on and allowed 3 h *ad lib* sleep prior to sacrifice for immunohistochemistry. ***Bottom:*** Confocal images of mCherry and cFos expression in the DG granule cell layer (white outlines) and hilar Sst+ interneurons. ***B)*** Expression of cFos in virally-transduced Sst+ interneurons was significantly higher in the hilus of hM3Dq-expressing mice than in mCherry control mice (*p* < 0.01, Student’s t-test).***C)*** Expression of cFos in surrounding DG neurons (in the DG granule cell layer) was reduced in hM3Dq-expressing mice (*p* < 0.05, Student’s t-test). **D) *Top:*** Experimental design: *SST-IRES-CRE* mice expressing mCherry (*n* = 4) or hM3Dq-mCherry (*n* = 5) in dorsal hippocampus underwent single-trial CFC at lights on, and then were immediately administered CNO (3 mg/kg, i.p.) and allowed *ad lib* sleep in their home cage. 24 h later, all mice were returned to the CFC context for assessment of contextual fear memory (CFM). ***Bottom:*** hM3Dq-expressing mice showed significant reductions in CFM consolidation (measured as % time freezing), compared with mCherry control mice (*p* < 0.001, Student’s t-test).

### Reducing cholinergic input to hippocampus improves sleep-dependent memory consolidation and increases hippocampal pS6 expression

Cholinergic input from MS selectively increases activity and structural plasticity in Sst+ interneurons, via muscarinic receptor activation (Gais and Born, 2004; Hajos et al., 1998; Lovett-Barron et al., 2014; Rasch et al., 2006; Raza et al., 2017; Schmid et al., 2016). Because acetylcholine release in the hippocampus is higher overall during wake vs. sleep (Teles-Grilo Ruivo et al., 2017), a reasonable assumption is that this drives higher Sst+ interneuron activity during SD. We tested whether altering medial septum (MS) cholinergic input to the hippocampus following CFC affected CFM consolidation. To do this, we transduced the MS of *Chat-CRE* mice with an AAV vector to express the inhibitory DREADD hM4Di in a Cre-dependent manner. Immunohistochemistry confirmed hM4Di-mCherry labeling of Chat+ terminals in the DG **(Figure 6A**), consistent with previous reports (Raza et al., 2017). To characterize the effects of reduced MS cholinergic input on network activity in the hippocampus, hM4Di-expressing mice were treated with CNO or VEH at lights on, and allowed 3 h *ad lib* sleep. Inhibition of cholinergic MS neurons in CNO-treated mice increased numbers of pS6+ neurons in DG relative to VEH-injected mice (*n* = 5 mice/group, *p* < 0.05, Student’s t-test) (**Figure 6B**). Transduced mice underwent single-trial CFC at lights on, after which they were immediately injected with either CNO or vehicle (VEH) (*n* = 10 mice/group), and were returned to their home cages for *ad lib* sleep. 24 h later, at lights on, mice were returned to the CFC context to assess CFM. Mice administered CNO showed significant increases in context-specific freezing compared to vehicle-treated mice (*p* < 0.05, Student’s t-test) (**Figure 6C**). These data suggest that MS cholinergic input to the hippocampus may mediate the state-dependent gating of hippocampal network activity in the same way that somatostatin does, thus acting a brake on memory consolidation mechanism.

**Figure 6.**
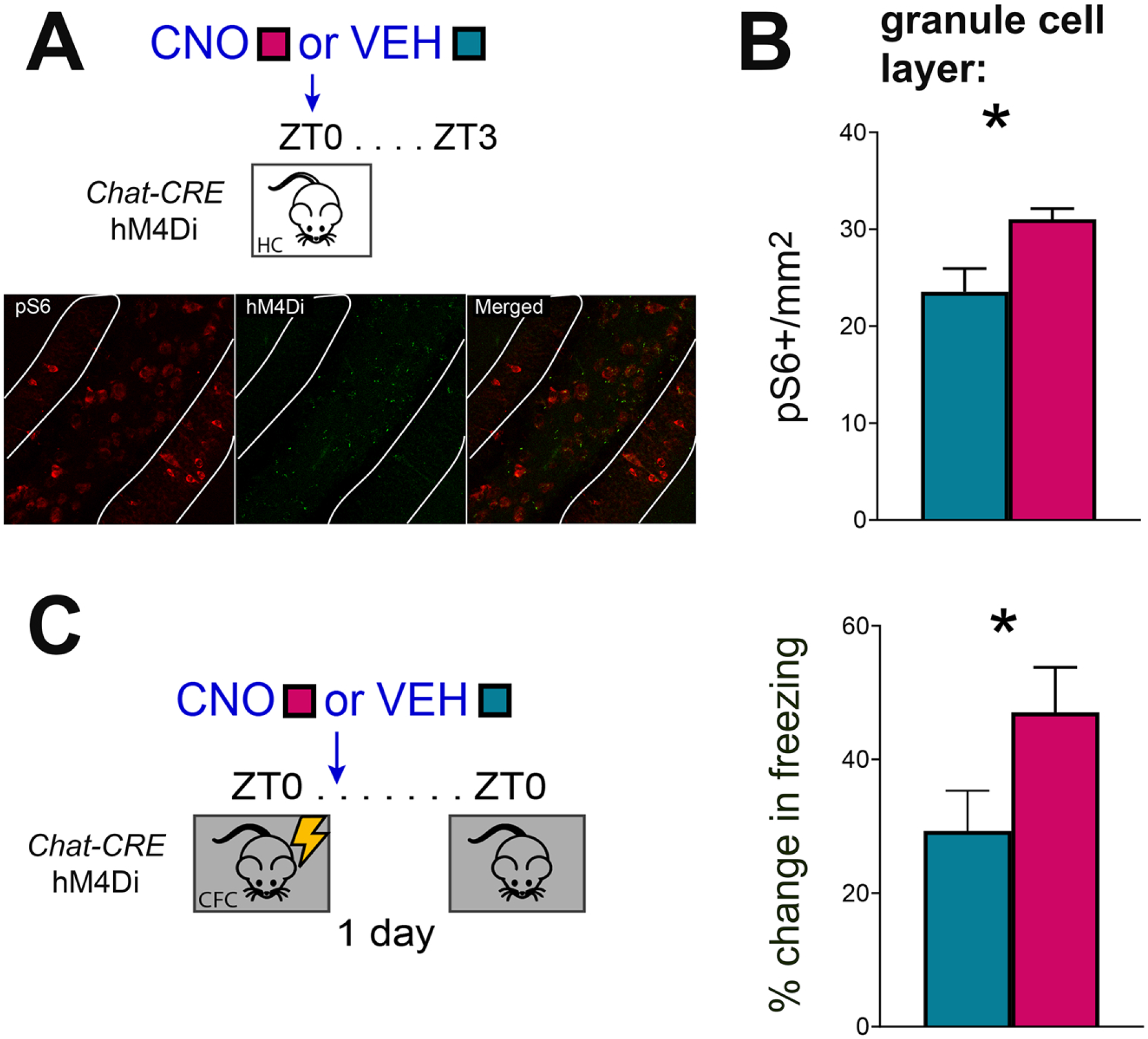
Reduced cholinergic input to the hippocampus increases DG network activity and improves sleep-associated memory consolidation. ***A) Top:*** Experimental design: *Chat-CRE* mice expressing hM4Di in the medial septum were administered CNO (3 mg/kg, i.p.) or vehicle (VEH) (*n* = 5/group) at ZT0, and were allowed 3 h *ad lib* sleep prior to sacrifice. ***Bottom:*** Representative images of reporter transgene expressing cholinergic terminals in DG and pS6+ expression in the DG granule cell layer (white lines) and hilus. ***B)*** Inhibition of cholinergic inputs to hippocampus increased numbers of pS6+ neurons in the DG granule cell layer. ***C) Right:*** Experimental design: *Chat-CRE* mice expressing hM4Di in the medial septum underwent single-trial CFC at lights on, after which they were immediately administered CNO (3 mg/kg, i.p.) or vehicle (VEH) (*n* = 10/group), and allowed *ad lib* sleep in their home cage. 24 h later, mice were returned to the CFC context for a CFM test.CNO-treated mice had higher numbers of pS6+ DG neurons after 3 h of *ad lib* sleep (*p*-value < 0.05, Student’s t-test). *Left:* CFM performance 24 h post-CFC was better in CNO-treated, relative to VEH-treated mice (*p*-value < 0.05, Student’s t-test).

## Discussion

Here we present converging lines of evidence that indicate SD disrupts activity in the dorsal hippocampus, and disrupts memory consolidation, via a Sst+ interneuron-mediated inhibitory gate. First, we find that, similar to *Arc* mRNA and Arc protein (Delorme et al., 2019), activity-dependent expression of pS6 in dorsal hippocampus increases across a brief period of sleep (**Figure S1**), but is reduced by a period of SD (**Figure 1**). This effect of SD is enhanced by prior learning (**Figure 1**, **Figure S1**), and seems to occur selectively in the hippocampus - it is not seen in the neocortex (**Figure S2**). We find that under SD conditions, pS6 expression in Sst+ interneurons (but not other cell types) is increased, rather than decreased (**Figure 4**). Using pS6 itself as an affinity tag, we took an unbiased TRAP-seq approach to characterize cell types active in the hippocampus during sleep vs. SD. We found that transcripts upregulated on pS6+ ribosomes after SD included those with expression unique to specific interneuron subtypes (*e.g*., *Sst*, *Npy*, and *Crhbp*), and markers of cholinergic and orexinergic neurons (*e.g*., *Cht*) (**Figure 2**, **Figure 3**). Hippocampal Sst+ interneurons are enriched in the same interneuron-specific markers identified as increasing in abundance after SD with pS6 TRAP-seq (**Figure 3B**). SD-driven changes in their expression appear to be caused by greater activity in these neurons as a function of SD (**Figure 3, Figure S3**).

Previous reports have shown that Sst+ interneurons gate DG network activity during memory acquisition, and that their activation likewise gates initial learning (Raza et al., 2017; Stefanelli et al., 2016). Considering we observe fewer activated DG neurons expressing either Arc or pS6 after SD (**Figure 1**), one possibility is that by driving higher firing activity in Sst+ interneurons, SD may disrupt memory consolidation by acting as an inhibitory gate – i.e., limiting activity in the surrounding network. Within the hippocampus, Sst+ interneurons target both neighboring pyramidal cells and other interneuron types (such as PV+ interneurons) for inhibition (Bloss et al., 2016; Harris et al., 2018; Katona et al., 1999; Katona et al., 2014; Pelkey et al., 2017; Somogyi et al., 2013). Our present data demonstrate that activation of Sst+ interneurons is sufficient to disrupt activity-regulated gene expression in neighboring neurons, and CFM consolidation (**Figure 5**). Because selective activation of Sst+ interneurons in the hippocampus is characteristic of SD, we conclude that this mechanism may explain the sleep-dependence of CFM consolidation (Graves et al., 2003; Ognjanovski et al., 2018; Vecsey et al., 2009). It may also explain other effects of SD on the dorsal hippocampal network, including disruption of long-term potentiation (LTP) (Havekes et al., 2016; Vecsey et al., 2009), reduction of plasticity-associated gene expression (Delorme et al., 2019), and decreases in dendritic spine density on pyramidal neurons (Havekes et al., 2016; Raven et al., 2019). CFM consolidation itself relies on intact network activity in dorsal hippocampus (Daumas et al., 2005), and is associated with increased network activity in structures such as CA1 (Ognjanovski et al., 2018; Ognjanovski et al., 2014), and regularization of spike-timing relationships during post-CFC sleep (Ognjanovski et al., 2018; Ognjanovski et al., 2014; Ognjanovski et al., 2017). Thus disruption of network activation via activation of an inhibitory circuit element during SD is likely to interfere with consolidation mechanisms. Critically, Sst+ interneurons may act as a gate on the hippocampal network, inhibiting sharp-wave ripple oscillations (Katona et al., 2014; Klausberger and Somogyi, 2008) and hippocampal-cortical communication (Abbas et al., 2018; Haam et al., 2018). Both of these features correlate with sleep-dependent consolidation of CFM (Ognjanovski et al., 2017; Xia et al., 2017).

Increased representation of cholinergic and orexinergic cell type-specific transcripts with SD using pS6-TRAP (**Figure 2**) is consistent with increased activity in lateral hypothalamic (orexinergic) and MS (cholinergic) inputs to dorsal hippocampus during wake (Kiyashchenko et al., 2002; Teles-Grilo Ruivo et al., 2017). Does activation of cholinergic or orexinergic inputs to the hippocampus likewise contribute to disruption of CFM consolidation? And do these modulators drive selective activation of hippocampal Sst+ interneurons during SD?

Behavioral data from both human subjects (Gais and Born, 2004; Rasch et al., 2006) and animal models (Inayat et al., 2020) demonstrate that reduced acetylcholine signaling is essential for the benefits of sleep for memory consolidation (Gais and Born, 2004; Rasch et al., 2006; Schmid et al., 2016). Our present data (**Figure 6**) are consistent with reductions in MS cholinergic input to the hippocampus being vital for sleep-dependent CFM consolidation. Available data suggest that these effects could be mediated by cholinergic regulation of Sst+ interneurons. Our present data demonstrate that pS6 expression in the hippocampus is augmented when MS cholinergic input is reduced (**Figure 6**), consistent with disinhibition. Others have found that stimulation of septohippocampal cholinergic neurons causes GABAergic inhibition of DG granule cells, mediated by cholinergic receptors on hilar (Sst- and Npy-expressing) interneurons (Pabst et al., 2016). In contrast, while less is known about the role of orexinergic signaling in memory consolidation, available data suggests that orexin can promote, rather than inhibit, consolidation (Mavanji et al., 2017). Moreover, orexinergic input to the hippocampus appears to activate glutamatergic neurons to a greater extent that GABAergic neurons (Stanley and Fadel, 2011).

An outstanding question is whether SD-associated, selective activation of Sst+ interneurons is driven mainly by network interactions (e.g., input from MS cholinergic interneurons) or cell-autonomous mechanisms (e.g. cell-type specific changes in expression of proteins that alter intrinsic excitability and neuronal firing). Our present data do not discriminate between these two mechanisms, but provide circumstantial evidence of both (**Figure 3** and **Figure 6**, respectively). A second outstanding question is whether these mechanisms are unique to the hippocampus, or whether similar selective activation of Sst+ interneurons is associated with SD in other structures, such as the neocortex. Recent findings from calcium imaging studies of mouse neocortex have demonstrated higher activity of Sst+ interneurons in superficial cortical layers during wake vs. NREM sleep (Niethard et al., 2017; Niethard et al., 2016). This effect (which would lead to reductions in dendrite-targeted inhibition during sleep) may be related to the recent finding of dendritic calcium spikes in these cortical layers during NREM oscillations (Seibt et al., 2017). Critically, activation of Sst+ interneurons in neocortex during active wake is driven by cholinergic signaling (Munoz et al., 2017). While this mechanism subserves effective circuit-level information processing during brief periods of arousal, one possibility is that extended wake may act as a gate, preventing information processing altogether - as we see evidence for here. Together, these findings suggest that a similar mechanism may underlie SD-induced disruption of memory consolidation mechanisms outside of the hippocampus (Puentes-Mestril and Aton, 2017), as well. Prior work has identified a number of intracellular pathways as being critical targets for SD-mediated disruption of memory in the hippocampus (Havekes et al., 2016; Tudor et al., 2016; Vecsey et al., 2009) and neocortex (Aton et al., 2009a; Dumoulin et al., 2015; Seibt et al., 2012). Our present data indicate that microcircuit regulation of brain activity is another SD-driven mechanism underlying the disruption of memory consolidation by sleep loss.

## Methods

### Mouse husbandry, handling, and behavioral procedures

All animal husbandry and experimental procedures were approved by the University of Michigan Institutional Animal Care and Use Committee (PHS Animal Welfare Assurance number D16-00072 [A3114-01]). Mice were maintained on a 12 h:12 h light:dark cycle with *ad lib* access to food and water. For behavioral experiments, 3-4 month old C57Bl6/J mice (Jackson) or transgenic mice on a C57Bl6/J background (see below) were individually housed with beneficial enrichment one week prior to experimental procedures, and were habituated to experimenter handling (5 min/day) for five days prior to experimental procedures. At lights on (ZT0), animals were either left in their home cage (HC) or underwent single-trial contextual fear conditioning (CFC). During CFC, mice were place in a novel conditioning chamber (Med Associates), and were allowed to explore the chamber freely for 2.5 min, after which they received a 2-s, 0.75 mA foot shock through the chamber’s grid floor. Mice remained in their conditioning chamber for an additional 28 s, after which they were then returned to their home cage. Mice were then either were permitted *ad lib* sleep (Sleep) or were sleep-deprived (SD) by gentle handling (Durkin et al., 2017; Ognjanovski et al., 2018) over the next 3-5 h.

### Translating Ribosome Affinity Purification (TRAP)

For pS6 RNA-sequencing (TRAP-seq) experiments, 3-4 month old C57Bl/6J mice were randomly assigned to one of four groups: HC + Sleep (*n* = 8), HC + SD (*n* = 7), CFC + Sleep (*n* = 8), CFC + SD (*n* = 8). Beginning at ZT3, animals were euthanized with an i.p injection of pentobarbital (Euthasol) and hippocampi were dissected in cold dissection buffer (1x HBSS, 2.5 mM HEPES [pH 7.4], 4 mM NaHCO_3_, 35 mM glucose, 100μg/ml cycloheximide). Hippocampal tissue was then transferred to a glass dounce homogenizer containing homogenization buffer (10 mM HEPES [pH 7.4], 150 mM KCl, 10 mM MgCl_2_, 2 mM DTT, cOmplete™ Protease Inhibitor Cocktail [Sigma-Aldrich, 11836170001], 100 U/mL RNasin® Ribonuclease Inhibitors [Promega, N2111], and 100 μg/mL cycloheximide) and manually homogenized on ice. Homogenate was transferred to 1.5 ml LoBind tubes (Eppendorf) and centrifuged at 4°C at 1000 g for 10 min. The resulting supernatant was transferred to a new tube, and 10% NP40 was added to the samples (90μL), and incubated 5 min on ice. Samples were centrifuged at 4°C at maximum speed for 10 min, 500μL supernatant transferred to a new LoBind tube, and incubated with anti-pS6 244-247 (ThermoFisher 44-923G; Knight et al., 2012). Antibody binding of the homogenate-antibody solution occurred over 1.5 h at 4°C with constant rotation. For affinity purification, 200 μl of Protein G Dynabeads (ThermoFisher, 10009D) were washed 3 times in 0.15M KCl IP buffer (10 mM HEPES [pH 7.4], 150 mM KCl, 10 mM MgCl_2_, 1% NP-40) and incubated in supplemented homogenization buffer (+10% NP-40). Following this step, supplemented buffer was removed, homogenate-antibody solution was added directly to the Dynabeads, and the solution was incubated for 1 h at 4°C with constant rotation. After incubation, the RNA-bound beads were washed four times in 900μL of 0.35M KCl (10mM HEPES [pH 7.4], 350 mM KCl, 10 mM MgCl_2_, 1% NP40, 2 mM DTT, 100 U/mL RNasin® Ribonuclease Inhibitors [Promega, N2111], and 100 μg/mL cycloheximide). During the final wash, beads were placed onto the magnet and moved to room temperature. After removing the supernatant, RNA was eluted by vortexing the beads vigorously in 350 μl RLT (Qiagen, 79216). Eluted RNA was purified using RNeasy Micro kit (Qiagen).

*Sst::RiboTag* mice were generated by crossing *SST-IRES-CRE* (B6N.Cg-Sst^tm2.1(SST-cre)Zjh^; Jackson) mice to the RiboTag^fl/fl^ (B6N.129-Rpl22^tm1.1Psam^/J; Jackson) mouse line to generate mice expressing HA-tagged Rpl22 protein in Sst+ interneurons. For Sst-TRAP, ribosomes and associated transcripts were affinity purified by incubating homogenate with 1/40 (10 μl) anti-HA antibody (Abcam, ab9110) (Shigeoka et al., 2018).

### RNA sequencing and data analysis

RNA-Seq was carried out at the University of Michigan’s DNA sequencing core. cDNA libraries were prepared by the core using Takara’s SMART-seq v4 Ultra Low Input RNA Kit (Takara 634888) and sequenced on Illumina’s NovaSeq 6000 platform. Sequencing reads (50 bp, paired end) were mapped to *Mus musculus* using Star v2.6.1a and quality checked with Multiqc(v1.6a0). Reads mapped to unique transcripts were counted with featureCounts (Liao et al., 2014). For weighted gene co-expression network analysis (WGCNA) analysis, raw counts for pS6 data was filtered to keep genes with at least 30 total reads across the 30 samples. The filtered reads were normalized using the DESeq2 variance stabilizing transformation (vst) function (Love et al., 2014) and filtered to keep genes with a variance larger than 0.03 among the 30 samples. The 1662 genes were retained and used for the network analysis in WGCNA (Langfelder and Horvath, 2008).

For cell type-specific expression analysis (CSEA), genes from the Brown and Magenta clusters were combined and uploaded into the CSEA Tool (http://genetics.wustl.edu/jdlab/csea-tool-2/), selecting Candidate Gene List from: Mice (Xu et al., 2014). Results for cell-types enriched in our pS6 clusters were analyzed at the most stringent specificity index (*p*SI < 0.0001). Observing significant values in cholinergic (+Chat), orexinergic (Hcrt+), and GABAergic (Pnoc+, Cort+) neurons, we plotted genes from the highest and second highest specificity index. To analyze how SD promotes pS6 enrichment of these cell type-specific transcripts, we calculated the Log_2_FC values of combined CFC and HC mice and assessed the effect of SD over sleep control mice (Love et al., 2014).

### Quantitative PCR (qPCR)

RNA from TRAP experiments was quantified by spectrophotometry (Nanodrop Lite, ThermoFisher). 50ng of RNA was reverse transcribed using iScript cDNA Synthesis (Bio-Rad, Catalog: 1708890) or SuperScript IV Vilo Master Mix (Invitrogen, Catalog: 11756060). qPCR was performed on diluted cDNA that employed either Power SYBR Green PCR Mix (Invitrogen 4367659) or TaqMan Fast Advanced Master Mix (Invitrogen, Catalog: 4444557). Primers were designed using Primer3(v. 0.4.0) and confirmed with NCBI primer Basic Local Alignment Search Tool (BLAST). qPCR reactions were measured using a CFX96 Real-Time System, in 96-well reaction plates (Bio-Rad). For pS6- and Sst-TRAP experiments, housekeeping genes for data normalization were determined by assessing the stability values prior to analysis (Andersen et al., 2004). Analyses compared *Pgk1*, *Gapdh*, *Actg1*, *Tuba4a*, *Tbp*, and *Hprt.* Results from both analyses independently found *Gapdh* and *Pgk1* to be the most stable and least altered housekeeping transcripts following SD or CFC. Therefore expression was normalized to the geometric mean of *Gapdh* and *Pgk1.* To assess differences in transcript abundance between groups, values were expressed as fold changes normalized to the mean values for mice in the HC + Sleep group. To measure relative enrichment of mRNA in pS6-TRAP or Sst-TRAP experiments, each sample was normalized to the geometric mean of *Pgk1* and *Gapdh* housekeeping transcripts and then normalized to the corresponding Input sample (TRAP Enrichment = 2^(ΔCt_target - ΔCt_housekeeping).

### Immunohistochemistry and protein expression analysis

Mice were injected with euthasol and perfused with cold 1xPBS followed by 4% paraformaldehyde. Brains were extracted and submerged in ice-cold fixative for 24hrs and transferred to 30% sucrose solubilized in 1xPBS. 50μm-thick coronal sections were cut on a cryostat. Tissue was blocked for 2-hours in 1% NGS and 0.3% Triton X-100 followed by 2-3 days of 4C° incubation in 1xPBS (5% NGS, 0.3% Triton X-100) with primary antibody(ies): pS6 S235-236 (Cell Signaling, Catalog: 4858, 1:500), pS6 S244-247(ThermoFisher, Catalog: 44-923G, 1:500), Sst (Millipore, MAB354, 1:200), Pvalb (Synaptic Systems, Catalog: 195004, 1:500), cFos (Abcam, Catalog: 190289, 1:500), Arc (Synaptic Systems, 156004, 1:500) by constant rotation. Sections were then washed 3x in 1xPBS (1% NGS, 0.2% Triton X-100) and incubated for 1hr in 1xPBS (5% NGS, 0.3% TX-100) and secondary antibody: Fluorescein (FITC) AffiniPure Goat Anti-Rabbit IgG (Jackson, Catalog: 111-095-003, 1:200), Donkey anti-Rat IgG Alexa Fluor 488 (ThermoFisher, A-21208), Goat Anti-Guinea Pig IgG Alexa Fluor 555 (Abcam, Catalog: ab150186, 1:200), Goat anti-Rabbit IgG Alexa Fluor 633 (ThermoFisher, Catalog: A-21070, 1:200). Tissue was then washed 3x in 1xPBS (1% NGS, 0.2% Triton X-100), 3x in 1xPBS, and then mounted on coverslips and embedded in ProLong Gold Antifade Mountant (ThermoFisher, Catalog: P10144).

For optical density (OD) calculations, 4 fluorescent microscope images were taken from each brain and analyzed in Fiji. A scorer blind to experimental condition collected optical density values from CA1 and CA3 pyramidal cell layers as well as background. For analysis, equally sized regions of interest (ROIs) were obtained for each image. OD values were background subtracted and normalized to HC + Sleep control groups. For DG cell counts, pS6 co-localization, and Sst and Pvalb quantification, images were captured using a 20x objective lens on a Leica SP5 laser scanning confocal microscope. Z-projected images were analyzed in MIPAR image analysis software in their raw grayscale format (Sosa et al., 2014) For DG cell counts, a non-local means filter was used to reduce image noise and an adaptive threshold applied to identify cell counts whose mean intensity values were 200% its surroundings. Colocalization and mean fluorescence intensity were determined by adaptive thresholding of fluorescent signals, quantifying percentage overlap of ROI obtained from both signals and mean fluorescent intensity values of underlying fluorescent signals.

### AAV virus injections, pharmacogenetic manipulations, and CFM testing

At age 3-4 months, male *SST-IRES-CRE* (B6N.Cg-Sst^tm2.1(SST-cre)Zjh^) or *Chat-CRE* (B6.FVB(Cg)-Tg(Chat-cre)GM53Gsat/Mmucd, MMRRC) mice underwent bilateral dorsal hippocampus or MS viral transduction. *SST-IRES-CRE* mice were transduced with either hM3dq-mCherry (pAAV-hSyn-DIO-hM3D(Gq)-mCherry, University of Pennsylvania Vector Core, Lot: V55836) or (as a control) an mCherry reporter (EF1A-DIO-mCherry, University of Pennsylvania Vector Core, Lot: PBK273-9). For both vectors, 1 μl of virus was injected using a 33-gauge beveled syringe needle into the dorsal hippocampus each hemisphere at a rate of 4 nL/s (2.1 mm posterior, 1.6 mm lateral, 2.1 mm ventral to Bregma). *Chat-*CRE mice were injected with hM4Di-mCitrine (AAV8 hSyn-DIO-HA-hM4D(Gi)-P2a-Citrine, University of Pennsylvania Vector Core, Lot: PBK399-9). 1 μl of virus was injected into the medial septum (0.75 mm anterior, 0.0 mm lateral, 4.0 mm ventral to Bregma).

After 2-4 weeks of postoperative recovery and daily handling as described above, mice underwent single-trial CFC at ZT0. Immediately after CFC, mice were injected i.p. with either 3 mg/kg clozapine N-oxide (CNO; Tocris, Catalog: 4936, Lot: 13D/233085) in 0.5% DMSO and saline, or 0.5% DMSO vehicle (VEH). All mice were then returned to their home cage for ad lib sleep, and were returned to the CFC context for CFM testing. Mice were video monitored and context-specific freezing behavior was quantified by a scorer blinded to experimental conditions as described previously (Ognjanovski et al., 2018; Ognjanovski et al., 2014; Ognjanovski et al., 2017). To verify effects of pharmacogenetic manipulations on the hippocampal network, 2 weeks following CFM tests, mice were administered CNO (or VEH) at lights on, and were allowed 3 h *ad lib sleep* prior to perfusion for immunohistochemical analysis of activity-dependent cFos or pS6 expression.

## Acknowledgements

The authors are grateful to members of the Aton lab for helpful feedback on this manuscript, and to Dr. Audrey Seasholtz for primers and feedback on the manuscript. This work was supported by research grants from the NIH (DP2 MH 104119) and the Human Frontiers Science Program (N023241-00_RG105) to SJA. We acknowledge support from the Bioinformatics Core of the University of Michigan Biomedical Research Core Facilities.

## Supplemental Information

**Figure S1.**
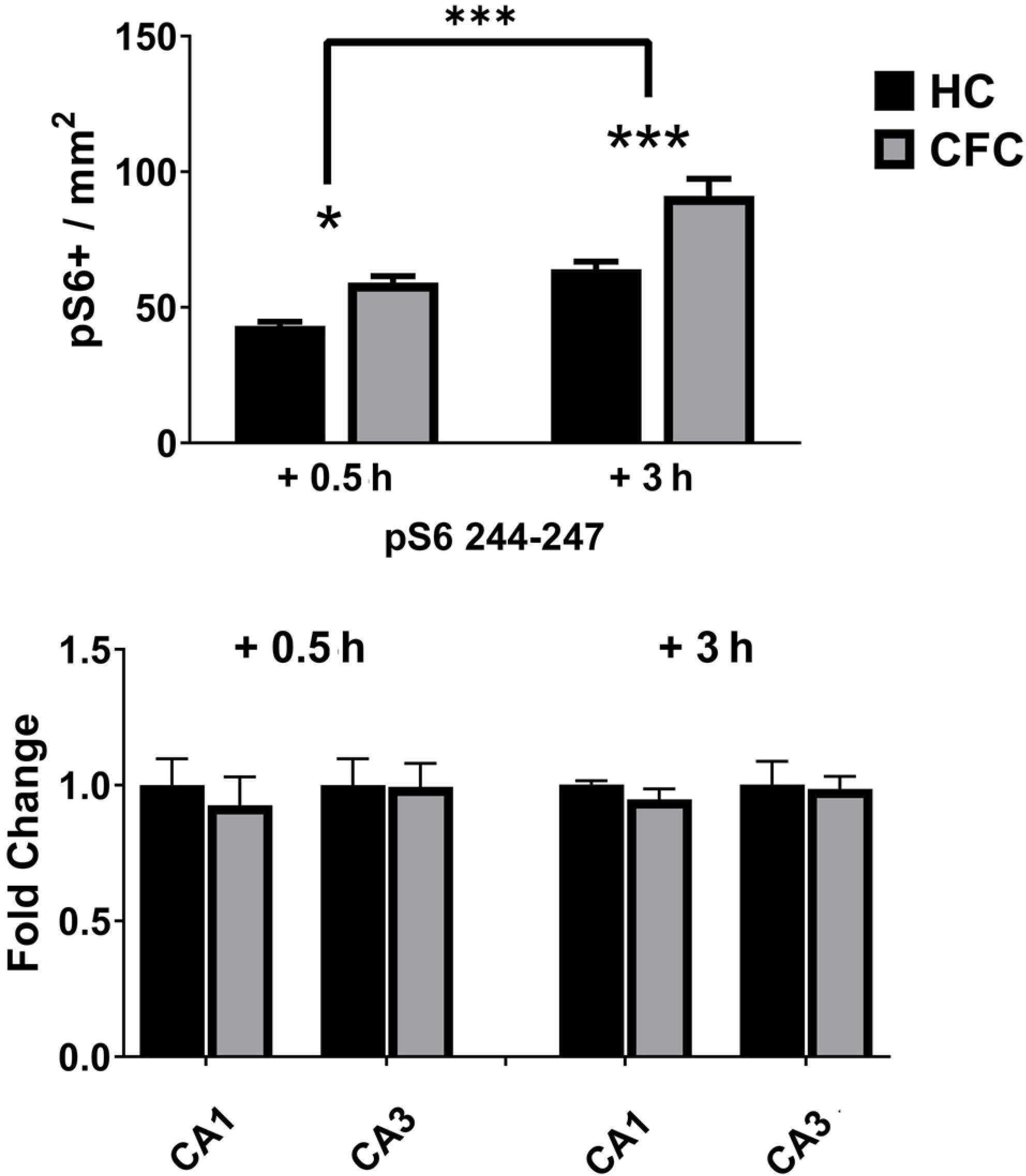
CFC-driven increases in hippocampal pS6 expression. ***Top:*** CFC (at ZT0) increased the number of pS6+ (S244-247) DG neurons at ZT0.5 h (*p* < 0.05, Holm-Sidak *post hoc* test) and ZT3 (*p* < 0.001, Holm-Sidak *post hoc* test). Both HC and CFC mice had greater numbers of pS6+ neurons at ZT3 (two-way ANOVA: main effect of time, *F* = 50.63, *p* < 0.001; main effect of CFC, *F* = 33.59, *p* < 0.001; time × CFC interaction, *N.S*.), likely reflecting the effect of sleep time between the time points. ***Bottom:*** pS6 expression was unchanged by CFC in CA1 and CA3 pyramidal regions.

**Figure S2.**
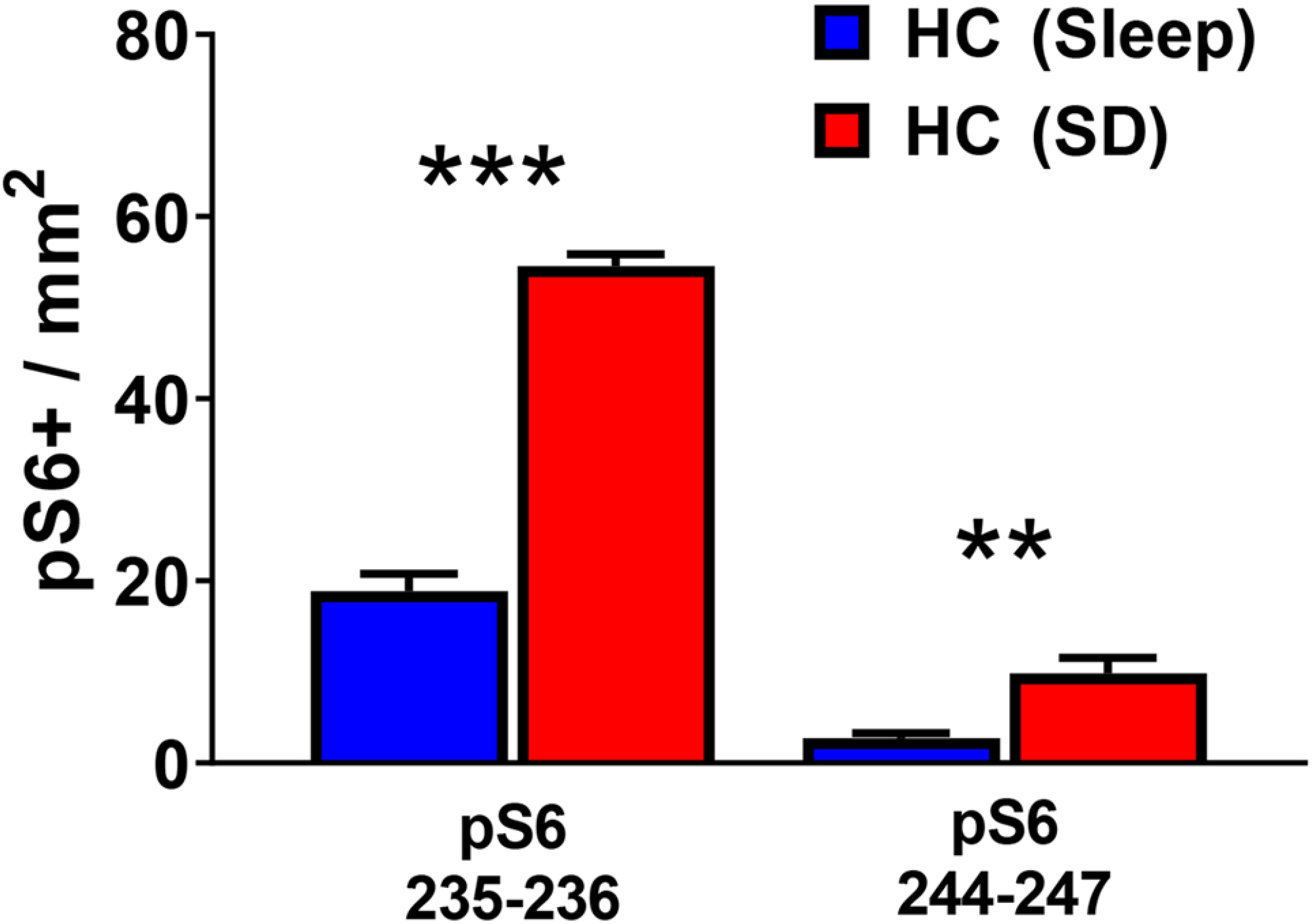
SD increases S6 phosphorylation in the neocortex. pS6 (S235-236 and S244-247) expression in neocortical regions dorsal hippocampus (i.e., primary somatosensory cortex). SD increased the numbers of pS6+ neurons using antibodies targeting pS6 S235-236 (*p* < 0.0001, Student’s t-test) and S244-247 (*p* < 0.01).

**Supplemental Figure S3.**
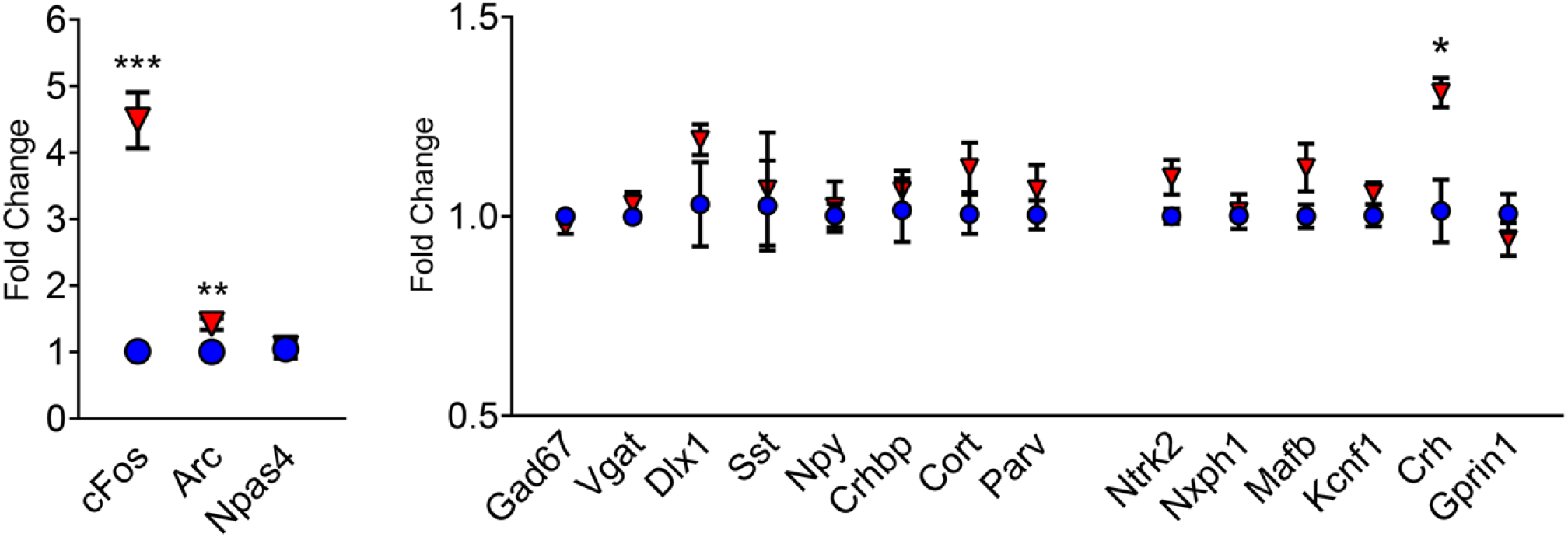
Whole hippocampus gene expression following 3 h SD or *ad lib* sleep. qPCR data for activity-dependent, GABAergic, and CSEA-predicted transcripts in whole hippocampus (Input) following 3h of SD (*n* = 4; red triangles) or Sleep (*n* = 5; blue circles). Sleep vs SD, ***, **, and * indicated *p* < 0.001, *p* < 0.01, and *p* < 0.05, respectively, Student’s t-test.

**Figure S4.**
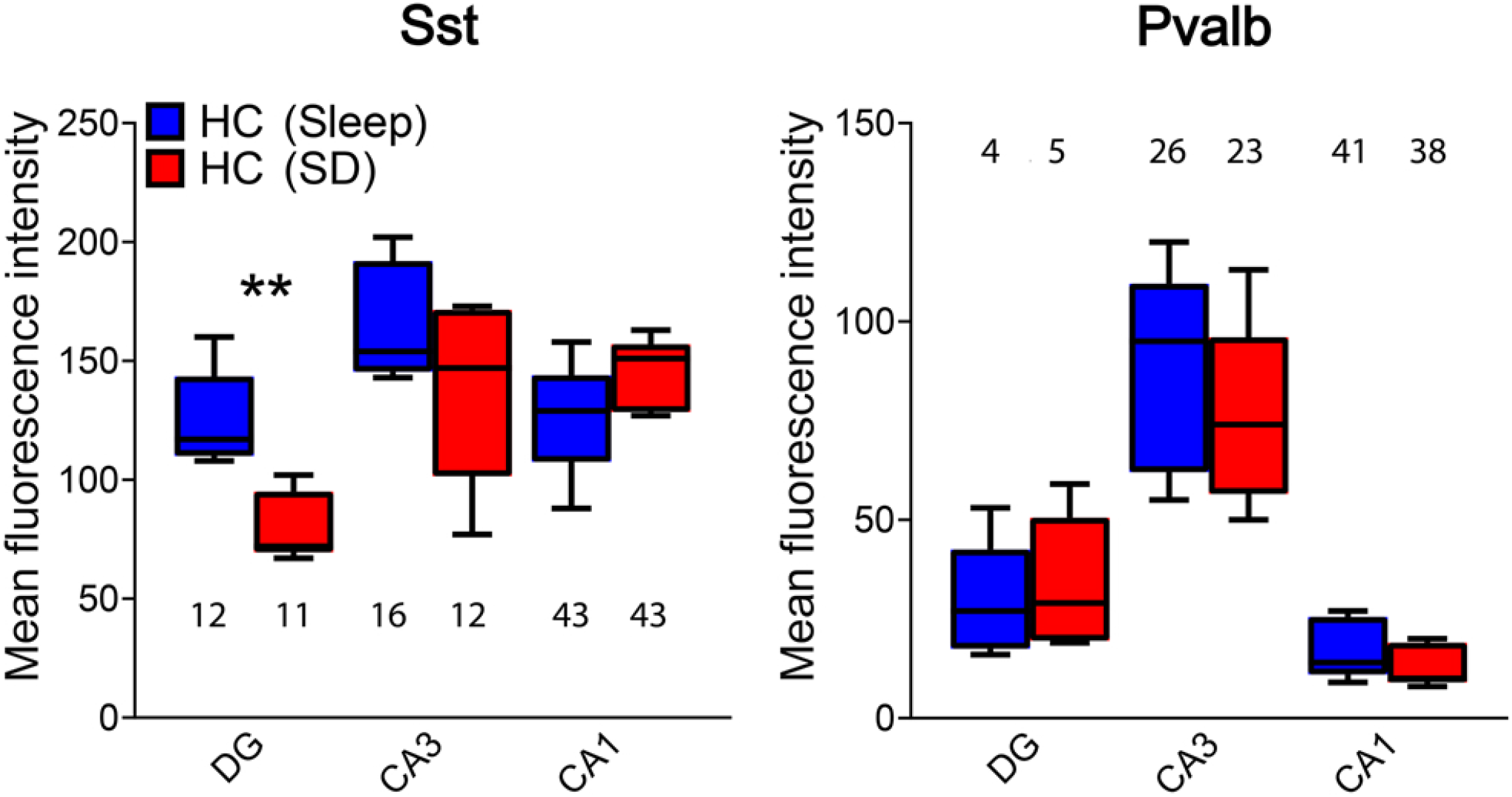
Mean Sst and Pvalb fluorescence intensity values in DG after sleep vs. SD. Mean fluorescence intensity values for Sst+ and Pvalb+ interneurons (numeric values indicate mean numbers of neurons quantified per animal) in DG following 3 h of *ad lib* sleep (*n* = 5 mice) or SD (*n* = 5 mice). 3-h SD reduced Sst staining intensity in DG neurons (Student’s t-test, *p* < 0.01).

**Supplemental Table 1 –** WGCNA cluster genes and CSEA analysis

## References cited

Abbas, A.I., Sundiang, M.J.M., Henoch, B., Morton, M.P., Bolkan, S.S., Park, A.J., Harris, A.Z., Kellendonk, C., and Gordon, J.A. (2018). Somatostatin Interneurons Facilitate Hippocampal-Prefrontal Synchrony and Prefrontal Spatial Encoding Neuron 100, 926–939.

Abel, T., Havekes, R., Salatin, J.M., and Walker, M.P. (2013). Sleep, Plasticity and Memory from Molecules to Whole-Brain Networks. Curr Biol 23, R774–788.

Andersen, C.L., Jensen, J.L., and Orntoft, T.F. (2004). Normalization of Real-Time Quantitative Reverse Transcription-PCR Data: A Model-Based Variance Estimation Approach to Identify Genes Suited for Normalization, Applied to Bladder and Colon Cancer Data Sets. Cancer Research 64, 5245–5250.

Aton, S.J., Seibt, J., Dumoulin, M., Jha, S.K., Steinmetz, N., Coleman, T., Naidoo, N., and Frank, M.G. (2009a). Mechanisms of sleep-dependent consolidation of cortical plasticity. Neuron 61, 454–466.

Aton, S.J., Seibt, J., and Frank, M.G. (2009b). Sleep and memory. In Encyclopedia of Life Science (Chichester: John Wiley and Sons, Ltd.).

Bloss, E.B., Cembrowski, M.S., Karsh, B., Colonell, J., Fetter, R.D., and Spruston, N. (2016). Structured Dendritic Inhibition Supports Branch-Selective Integration in CA1 Pyramidal Cells. Neuron 89, 1016–1030.

Bomkamp, C., Tripathy, S.J., Bengtsson Gonzalez, C., Hjerling-Leffler, J., Craig, A.M., and Pavlidis, P. (2019). Transcriptomic correlates of electrophysiological and morphological diversity within and across excitatory and inhibitory neuron classes. PLoS Computational Biology 15.

Daumas, S., Halley, H., Frances, B., and Lassalle, J.M. (2005). Encoding, consolidation, and retrieval of contextual memory: differential involvement of dorsal CA3 and CA1 hippocampal subregions. Learn Mem 12, 375–382.

Delorme, J.E., Kodoth, V., and Aton, S.J. (2019). Sleep loss disrupts Arc expression in dentate gyrus neurons. Neurobiol Learn Mem 160, 73–82.

Doyle, J.P., Dougherty, J.D., Heiman, M., Schmidt, E.F., Stevens, T.R., Ma, G., Bupp, S., Shrestha, P., Shah, R.D., Doughty, M.L., et al. (2008). Application of a Translational Profiling Approach for the Comparative Analysis of CNS Cell Types. Cell 135, 749–762.

Dumoulin, M.C., Aton, S.J., Watson, A.J., Renouard, L., Coleman, T., and Frank, M.G. (2015). Extracellular Signal-Regulated Kinase (ERK) Activity During Sleep Consolidates Cortical Plasticity In Vivo. Cereb Cortex 25, 507–515.

Durkin, J., Suresh, A.K., Colbath, J., Broussard, C., Wu, J., Zochowski, M., and Aton, S.J. (2017). Cortically coordinated NREM thalamocortical oscillations play an essential, instructive role in visual system plasticity. Proceedings National Academy of Sciences 114, 10485–10490.

Gais, S., and Born, J. (2004). Low acetylcholine during slow-wave sleep is critical for declarative memory consolidation. PNAS 101, 2140–2144.

Graves, L.A., Heller, E.A., Pack, A.I., and Abel, T. (2003). Sleep deprivation selectively impairs memory consolidation for contextual fear conditioning. Learn Mem 10, 168–176.

Haam, J., Zhou, J., Cui, G., and Yakel, J.L. (2018). Septal cholinergic neurons gate hippocampal output to entorhinal cortex via oriens lacunosum moleculare interneurons. Proc Natl Acad Sci USA 115, E1886–1895.

Hajos, N., E.C., P., Acsady, L., Levey, A.I., and Freund, T.F. (1998). Distinct Interneuron Types Express m2 Muscarinic Receptor Immunoreactivity on Their Dendrites or Axon Terminals in the Hippocampus Neuroscience 82, 355–376.

Harris, K.D., Hochgerner, H., Skene, N.G., Magno, L., Katona, L., Bengtsson Gonzales, C., Somogyi, P., Kessaris, N., Linnarsson, S., and Hjerling-Leffler, J. (2018). Classes and continua of hippocampal CA1 inhibitory neurons revealed by single-cell transcriptomics. PLoS Biology 16, e2006387.

Havekes, R., and Abel, T. (2017). The tired hippocampus: the molecular impact of sleep deprivation on hippocampal function. Curr Opin Neurobiol 44, 13–19.

Havekes, R., Park, A.J., Tudor, J.C., Luczak, V.G., Hansen, R.T., Ferri, S.L., Bruinenberg, V.M., Poplawski, S.G., Day, J.P., Aton, S.J., et al. (2016). Sleep deprivation causes memory deficits by negatively impacting neuronal connectivity in hippocampal area CA1. eLife 5, pii: e13424.

Inayat, S., Qandeel, Nasariahangarkoaee, M., Singh, S., McNaughton, B.L., Whishaw, I.Q., and Mohajerani, M.H. (2020). Low acetylcholine during early sleep is important for motor memory consolidation. Sleep 43.

Katona, I., Acsady, L., and Freund, T.F. (1999). Postsynaptic targets of somatostatin-immunoreactive interneurons in the rat hippocampus. Neuroscience 88, 37–55.

Katona, L., Lapray, D., Viney, T.J., Oulhaj, A., Borhegyi, Z., Micklem, B.R., Klausberger, T., and Somogyi, P. (2014). Sleep and Movement Differentiates Actions of Two Types of Somatostatin-Expressing GABAergic Interneuron in Rat Hippocampus Neuron 82, 872–886.

Kiyashchenko, L.I., Mileykovskiy, B.Y., Maidment, N., Lam, H.A., Wu, M.-F., John, J., Peever, J., and Siegel, J.M. (2002). Release of Hypocretin (Orexin) during Waking and Sleep States. JNeurosci 22, 5282–5286.

Klausberger, T., and Somogyi, P. (2008). Neuronal Diversity and Temporal Dynamics: The Unity of Hippocampal Circuit Operations Science 321, 53–57.

Knight, Z.A., Tan, K., Birsoy, K., Schmidt, S., Garrison, J.L., Wysocki, R.W., Emiliano, A., Ekstrand, M.I., and Friedman, J.M. (2012). Molecular profiling of activated neurons by phosphorylated ribosome capture. Cell 151, 1126–1137.

Kosaka, T., Wu, J.-Y., and Benoit, R. (1998). GABAergic neurons containing somatostatin-like immunoreactivity in the rat hippocampus and dentate gyrus. Experimental Brain Res 71, 388–398.

Langfelder, P., and Horvath, S. (2008). WGCNA: an R package for weighted correlation network analysis. BMC Bioinformatics 9.

Liao, Y., Smyth, G.K., and Shi, W. (2014). featureCounts: An Efficient General Purpose Program for Assigning Sequence Reads to Genomic Features Bioinformatics 30, 329–330.

Love, M.I., Huber, W., and Anders, S. (2014). Moderated estimation of fold change and dispersion for RNA-seq data with DESeq2. Genome Biology 15.

Lovett-Barron, M., Kaifosh, P., Kheirbek, M.A., Danielson, N., Zaremba, J.D., Reardon, T.R., Turi, G.F., Hen, R., Zemelman, B.V., and Losonczy, A. (2014). Dendritic Inhibition in the Hippocampus Supports Fear Learning Science 343, 857–863.

Mavanji, V., Butterick, T.A., Duffy, C.M., Nixon, J.P., Billington, C.J., and Kotz, C.M. (2017). Orexin/hypocretin treatment restores hippocampal-dependent memory in orexin-deficient mice. Neurobiol Learn Mem 146, 21–30.

Munoz, W., Tremblay, R., Levenstein, D., and Rudy, B. (2017). Layer-specific modulation of neocortical dendritic inhibition during active wakefulness. Science 355, 954–959.

Niethard, N., Burgalossi, A., and Born, J. (2017). Plasticity during Sleep Is Linked to Specific Regulation of Cortical Circuit Activity. Front Neural Circuits 11.

Niethard, N., Hasegawa, M., Itokazu, T., Oyanedel, C.N., Born, J., and Sato, T.R. (2016). Sleep-Stage-Specific Regulation of Cortical Excitation and Inhibition. Curr Biol 26, 2739–2749.

Ognjanovski, N., Broussard, C., Zochowski, M., and Aton, S.J. (2018). Hippocampal Network Oscillations Rescue Memory Consolidation Deficits Caused by Sleep Loss. Cereb Cortex 28, 3711–3723.

Ognjanovski, N., Maruyama, D., Lashner, N., Zochowski, M., and Aton, S.J. (2014). CA1 hippocampal network activity changes during sleep-dependent memory consolidation. Front Syst Neurosci 8, 61.

Ognjanovski, N., Schaeffer, S., Mofakham, S., Wu, J., Maruyama, D., Zochowski, M., and Aton, S.J. (2017). Parvalbumin-expressing interneurons coordinate hippocampal network dynamics required for memory consolidation. Nature Communications 8, 15039.

Pabst, M., Braganza, O., Dannerberg, H., Hu, W., Pothmann, L., Rosen, J., Mody, I., van Loo, K., Deisseroth, K., Becker, A.J., et al. (2016). Astrocyte Intermediaries of Septal Cholinergic Modulation in the Hippocampus. Neuron 90, 853–865.

Pelkey, K.A., Chittajallu, R., Craig, M.T., Tricoire, L., Wester, J.C., and McBain, C.J. (2017). Hippocampal GABAergic Inhibitory Interneurons. Physiol Rev 97, 1619–1747.

Pirbhoy, P.S., Farris, S., and Steward, O. (2016). Synaptic activation of ribosomal protein S6 phosphorylation occurs locally in activated dendritic domains. Learn Mem 23, 255–269.

Prince, T.M., Wimmer, M., Choi, J., Havekes, R., Aton, S., and Abel, T. (2014). Sleep deprivation during a specific 3-hour time window post-training impairs hippocampal synaptic plasticity and memory. Neurobiol Learn Mem 109, 122–130.

Puentes-Mestril, C., and Aton, S.J. (2017). Linking network activity to synaptic plasticity during sleep: hypotheses and recent data. Frontiers in Neural Circuits 11, doi: 10.3389/fncir.2017.00061.

Rasch, B.H., Born, J., and Gais, S. (2006). Combined blockade of cholinergic receptors shifts the brain from stimulus encoding to memory consolidation. Journal of Cognitive Neuroscience 18, 793–802.

Raven, F., Meerlo, P., Van der Zee, E.A., Abel, T., and Havekes, R. (2019). A brief period of sleep deprivation causes spine loss in the dentate gyrus of mice. Neurobiol Learn Mem 160, 83–90.

Raza, S.A., Albrecht, A., Caliskan, G., Muller, B., Demiray, Y.E., Ludewig, S., Meis, S., Faber, N., Hartig, R., Schraven, B., et al. (2017). HIPP neurons in the dentate gyrus mediate the cholinergic modulation of background context memory salience. Nat Communications 8.

Sanz, E., Bean, J.C., Carey, D.P., Quintana, A., and McKnight, G.S. (2019). RiboTag: Ribosomal Tagging Strategy to Analyze Cell-Type-Specific mRNA Expression In Vivo. Curr Protoc Neurosci 88, e77.

Schmid, L.C., Mittag, M., Poll, S., Steffen, J., Wagner, J., Geis, H.-R., Schwarz, I., Schmidt, B., Schwarz, M.K., Remy, S., et al. (2016). Dysfunction of Somatostatin-Positive Interneurons Associated With Memory Deficits in an Alzheimer’s Disease Model Neuron 92, 114–125.

Seibt, J., Dumoulin, M., Aton, S.J., Coleman, T., Watson, A., Naidoo, N., and Frank, M.G. (2012). Protein synthesis during sleep consolidates cortical plasticity in vivo. Curr Biol 22, 676–682.

Seibt, J., Richard, C.J., Sigl-Glockner, U., Takahashi, N., Kaplan, D.I., Doron, G., de Limoges, D., Bocklisch, C., and Larkum, M.E. (2017). Cortical dendritic activity correlates with spindle-rich oscillations during sleep in rodents. Nat Commun 8, 1838.

Shigeoka, T., Jung, J., Holt, C.E., and Jung, H. (2018). Axon-TRAP-RiboTag: Affinity Purification of Translated mRNAs from Neuronal Axons in Mouse In Vivo. In RNA Detection (Methods in Molecular Biology) (New York: Humana Press), pp. 85–94.

Somogyi, P., Katona, L., Klausberger, T., Lasztoczi, B., and Viney, T.J. (2013). Temporal redistribution of inhibition over neuronal subcellular domains underlies state-dependent rhythmic change of excitability in the hippocampus. Philos Trans R Soc Lond B Biol Sci 369, 20120518.

Sosa, J.M., Huber, D.E., Welk, B., and H.L., F. (2014). Development and application of MIPAR™: a novel software package for two- and three-dimensional microstructural characterization. Integrating Materials and Manufacturing Innovation 3, 123–140.

Stanley, E.M., and Fadel, J.R. (2011). Aging-related alterations in orexin/hypocretin modulation of septo-hippocampal amino acid neurotransmission. Neuroscience 195, 70–90.

Stefanelli, T., Bertollini, C., Luscher, C., Muller, D., and Mendez, P. (2016). Hippocampal Somatostatin Interneurons Control the Size of Neuronal Memory Ensembles. Neuron 89, 1074–1085.

Taniguchi, H., He, M., Wu, P., Kim, S., Paik, R., Sugino, K., Kvitsani, D., Fu, Y., Lu, J., Lin, Y., et al. (2011). A Resource of Cre Driver Lines for Genetic Targeting of GABAergic Neurons in Cerebral Cortex. Neuron 71, 995–1013.

Teles-Grilo Ruivo, L.M., Baker, K.L., Conway, M.L., Isaac, J.T.R., Lowry, J.P., and Mellor, J.R. (2017). Coordinated Acetylcholine Release in PrefrontalCortex and Hippocampus Is Associated with Arousal and Reward on Distinct Timescales. Cell Reports 18, 905–917.

Tudor, J.C., Davis, E.J., Peixoto, L., Wimmer, M.E., van Tilborg, E., Park, A.J., Poplawski, S.G., Chung, C.W., Havekes, R., Huang, J., et al. (2016). Sleep deprivation impairs memory by attenuating mTORC1-dependent protein synthesis. Sci Signal 9, ra41.

Vecsey, C.G., Baillie, G.S., Jaganath, D., Havekes, R., Daniels, A., Wimmer, M., Huang, T., Brown, K.M., Li, X.Y., Descalzi, G., et al. (2009). Sleep deprivation impairs cAMP signalling in the hippocampus. Nature 461, 1122–1125.

Vecsey, C.G., Peixoto, L., Choi, J.H., Wimmer, M., Jaganath, D., Hernandez, P.J., Blackwell, J., Meda, K., Park, A.J., Hannenhalli, S., et al. (2012). Genomic analysis of sleep deprivation reveals translational regulation in the hippocampus. Physiol Genomics 44, 981–991.

Xia, F., Richards, B.A., Tran, M.M., Josselyn, S.A., Takehara-Nishiuchi, K., and Frankland, P.W. (2017). Parvalbumin-positive interneurons mediate neocortical-hippocampal interactions that are necessary for memory consolidation. eLife 6, e27868.

Xu, X., Wells, A.B., O’Brien, D.R., Nehorai, A., and Dougherty, J.D. (2014). Cell Type-Specific Expression Analysis to Identify Putative Cellular Mechanisms for Neurogenetic Disorders. J Neurosci 34, 1420–1431.

